# Homoeolog expression in polyploid wheat mutants shows limited transcriptional compensation

**DOI:** 10.1101/2025.07.01.662569

**Authors:** Delfi Dorussen, Emilie Knight, James Simmonds, Philippa Borrill

## Abstract

Active transcriptional compensation between gene duplicates (paralogs or homoeologs) has been proposed to facilitate functional redundancy, whereby mutations in multiple gene copies are required before a phenotype is observed. We tested whether transcriptional compensation occurs between homoeologs in response to premature termination codon (PTC) mutations in mutagenised wheat lines. Only ∼3% of cases showed homoeologous upregulation in response to PTC mutations, suggesting that there is no widespread active transcriptional compensation between wheat homoeologs.

## Introduction

Whole genome duplications (WGDs), hybridisations, and small-scale duplications have resulted in an abundance of gene duplicates in plant genomes – on average, 65% of genes in plant genomes have a paralog (Panchy *et al*., 2016). These duplications can provide opportunities for genetic innovation, when mutations cause one of the copies to adopt a novel function (neo-functionalisation) (Birchler & Yang, 2022). Alternatively, the presence of gene duplicates can result in functional redundancy, whereby the effect of a loss-of-function mutation in one gene copy is masked by the remaining functional copy. It is debated whether this phenotypic robustness is evolutionarily advantageous, or whether selection for lower expression noise results in retention of duplicated genes (Pires & Conant, 2016; Iohannes & Jackson, 2023). Nevertheless, this buffering of deleterious mutations has hindered the functional characterisation of genes (for example, through gene knockouts) and reduced the variation available for crop breeding (Uauy *et al*., 2017).

Hybridisation and polyploidisation events are widespread in the evolutionary history of the angiosperms and have given rise to major cereal crops such as bread wheat (*Triticum aestivum*) and pasta wheat (*Triticum turgidum* ssp. *durum*) (Matsuoka, 2011). Bread wheat is a hexaploid, formed by the merger of three diploid progenitor species. The hexaploid wheat genome therefore consists of three subgenomes (A, B, and D) (El Baidouri *et al*., 2017; IWGSC *et al*., 2018). Pasta wheat is a tetraploid and its genome consists of the A and B subgenomes (El Baidouri *et al*., 2017). As the wheat progenitor species were closely related, many genes are present in highly similar copies across the subgenomes – these duplicate genes formed by polyploidisation are known as homoeologs and share an average nucleotide sequence identity of 97.2% (Schreiber *et al*., 2012; IWGSC *et al*., 2018). Similar to paralogs, functional redundancy can exist between homoeologs, and loss-of-function mutations in multiple homoeologs may be required before a phenotype is observed. For example, in hexaploid wheat, loss-of-function mutations in all three homoeologs of the *Ms26* gene are required to confer male sterility, while single mutants have no reduction in fertility (Singh *et al*., 2017). Similarly, only triple mutants in *Qsd1* have extended seed dormancy (Abe *et al*., 2019). This redundancy may result in hidden variation in single homoeologs that cannot be observed until mutations in multiple homoeologs are combined (Borrill *et al*., 2015).

However, it remains unclear how phenotypic compensation between gene duplicates occurs. Active transcriptional compensation, the transcriptional upregulation of genes with a high degree of sequence similarity (such as paralogs or homoeologs) in response to a loss-of-function mutation in a gene, has been proposed as an explanation for functional redundancy.

Such transcriptional compensation has been observed in multiple species, including *Caenorhabditis elegans* (Serobyan *et al*., 2020) and *Danio rerio* (zebrafish; (El-Brolosy *et al*., 2019; Ma *et al*., 2019)). The degree of transcriptional compensation between paralogs is also hypothesised to underly the penetrance of mutations – for example, in zebrafish, a greater degree of craniofacial distortion was observed due to mutation of *MEF2CA* in the absence of upregulation of the *MEF2C* paralogs (Bailon-Zambrano *et al*., 2022).

Transcriptional compensation between paralogs has also been documented in numerous plant species, for example in the *CLE* gene family. Transcriptional upregulation of *CLE9* (a *CLV3* paralog) is observed in *clv3* mutants in *Solanum lycopersicum* (tomato), *Petunia hybrida* (petunia) and *Physalis grisea* (groundcherry) (Rodriguez-Leal *et al*., 2019; Kwon *et al*., 2022). As such, *clv3* mutants in these species have weaker phenotypes than the *clv3* mutant in *Nicotiana benthamiana* (tobacco), in which *CLE9* has become pseudogenised (Kwon *et al*., 2022). However, active compensation is not always observed between paralogs in plants – for example, in *Arabidopsis thaliana*, there was no transcriptional upregulation of *RPL23aA* in response to knock-out of its paralog *RPL23aB*, or vice versa (Degenhardt & Bonham-Smith, 2008; Xiong, W *et al*., 2020). It is unknown whether active transcriptional compensation occurs in wheat or other polyploid plants, buffering the effects of mutations in individual homoeologs.

Here we assessed whether transcriptional compensation is prevalent between homoeologs in hexaploid and tetraploid wheat using ethyl methanesulfonate (EMS) mutagenised TILLING (Targeting Induced Local Lesions in Genomes) lines (Krasileva *et al*., 2017). Each wheat TILLING line has a large number of mutations (more than 5,000 in the hexaploid cultivar Cadenza) (Krasileva *et al*., 2017), allowing us to simultaneously screen the effect of many mutations. We performed RNA-sequencing and differential gene expression analysis to determine whether genes are frequently upregulated in response to a loss-of-function mutation in one of their homoeologs. We found no evidence for widespread active transcriptional compensation between homoeologs in hexaploid or tetraploid wheat, indicating that such a mechanism is unlikely to be the primary cause of functional redundancy between homoeologs in polyploid wheat.

## Results

To test whether active transcriptional compensation occurs between homoeologs in hexaploid wheat, we screened four representative EMS-mutagenised TILLING lines (cv. Cadenza) to identify homoeolog groups with a premature termination codon (PTC) mutation in one of the homoeologs. PTC mutations are expected to cause truncation of the protein encoded by the gene, thus resulting in loss of function that could be associated with up-regulation of the gene’s homoeologs to provide a buffering effect. Differential expression of the gene affected by the PTC mutation and its homoeologs in the TILLING line relative to wild type (WT) was determined by analysis of RNA-sequencing data (**Figure 1a**).

**Figure 1.**
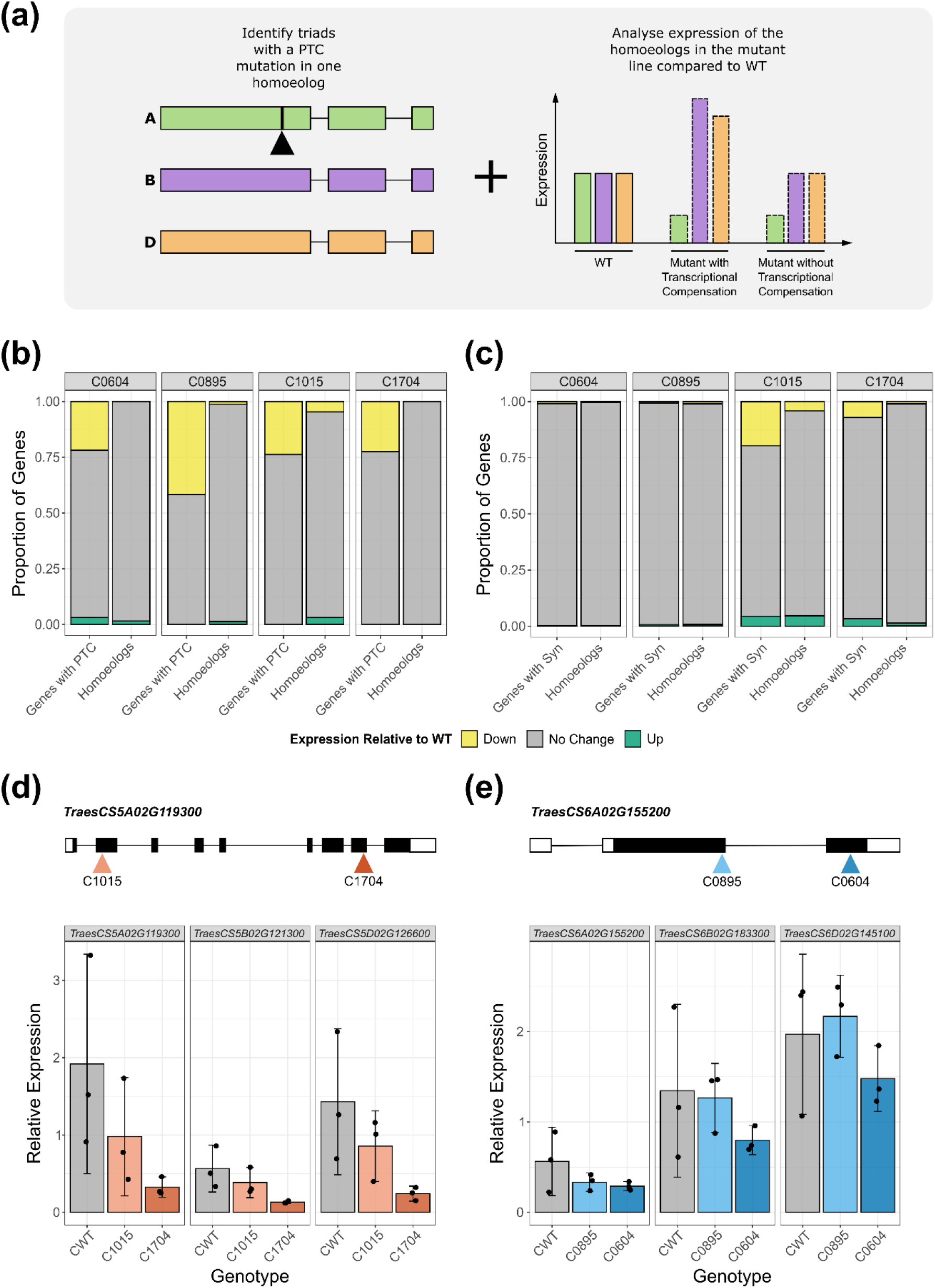
Absence of active transcriptional compensation between homoeologs in response to premature termination codon (PTC) mutations in hexaploid wheat. (a) Homoeolog groups with a PTC mutation in one homoeolog were identified in mutagenised wheat lines, and homoeolog-specific expression was analysed relative to the WT control. This allowed us to identify homoeolog groups with and without transcriptional compensation. (b) Expression of genes with a premature termination codon (PTC) mutation (left bar) and their homoeologs (right bar) in each of the mutagenised Cadenza lines. (c) Expression of genes with a synonymous mutation (left bar) and their homoeologs (right bar) in each of the mutagenised lines. In (b) and (c) the proportion of genes that are down-regulated (FDR adjusted p-value < 0.05) is shown in yellow, up-regulated (FDR adjusted p-value < 0.05) in green, and no change in expression in grey, all relative to Cadenza WT. (d) The location of PTC mutations in *TraesCS5A02G119300* in the C1015 and C1704 lines and the relative expression of *TraesCS5A02G119300* and its homoeologs (*TraesCS5B02G121300* and *TraesCS5D02G126600*) in Cadenza WT (CWT) (grey), C1015 (light orange), and C1704 (dark orange). (e) The location of PTC mutations in *TraesCS6A02G155200* in the C0895 and C0604 lines and the relative expression of *TraesCS6A02G155200* and its homoeologs (*TraesCS6B02G183300* and *TraesCS6D02G145100*) in Cadenza WT (CWT) (grey), C0895 (light blue), and C0604 (dark blue). In (d) and (e) in the gene schematics, black rectangles represent exons, white rectangles represent untranslated regions, and lines represent introns. Error bars represent the 95% confidence interval around the mean, estimated as 1.96 x standard error.

Across the four mutagenised lines (C0604, C0895, C1015, and C1704), we identified 158 unique homoeolog groups with a PTC mutation in one homoeolog. In 20.6% (C0604) to 41.7% (C0895) of cases, the homoeolog with the PTC mutation was down-regulated in the mutagenised line relative to WT (FDR adjusted p-value < 0.05), indicating that the abundance of functional transcript was reduced (**Figure 1b**). However, despite our relaxed threshold for detection, we did not observe widespread up-regulation of the homoeologous genes – across the mutagenised lines, we only identified 4 homoeolog groups (2.5%) in which at least one of the homoeologs was up-regulated (FDR adjusted p-value < 0.05, **Figure 1b, Supplementary Table 1**).

Furthermore, the proportion of homoeolog groups with up-regulated homoeologs was not significantly different for homoeolog groups affected by PTC mutations or synonymous mutations (2.5% of homoeolog groups with a PTC mutation, 3.8% of homoeolog groups with a synonymous mutation; χ^2^ test p-value = 0.401, **Figure 1c**). Active transcriptional compensation is not expected to occur in response to synonymous mutations as they are not expected to affect protein function. This suggests that active transcriptional compensation is not the default mechanism for buffering single homoeolog mutations.

We found a similar lack of active transcriptional compensation in an independent RNA-sequencing dataset of two EMS-mutagenised lines produced by Xiong, HC *et al*. (2020). Thirty homoeolog groups with a PTC mutation were identified, of which only one (3.3%) showed up-regulation of the homoeologs (**Supplementary Figure 1a, Supplementary Table 1**). Again, there was no significant difference between the proportion of up-regulated homoeologs in groups affected by a PTC mutation compared to those affected by a synonymous mutation (3.5% of homoeolog groups with a synonymous mutation, χ^2^ test p-value = 0.951, **Supplementary Figure 1b**).

Next, we investigated whether the location of the PTC mutation within the transcript affects whether active transcriptional compensation occurs. We hypothesised that PTC mutations occurring earlier within the coding sequence would have a greater impact on protein function and therefore promote more compensatory up-regulation. We identified genes with multiple PTC mutations across the four Cadenza mutagenised lines, differing in their position within the gene. Accordingly, we found *TraesCS5A02G119300* (with an early PTC in C1015 and a late PTC in C1704) and *TraesCS6A02G155200* (with an early PTC in C0895 and a late PTC in C0604) (**Figure 1d-e**). The relative expression of *TraesCS5A02G119300, TraesCS6A02G155200* and their homoeologs in the mutagenised lines compared to WT was determined by RT-qPCR. We found no significant effect of genotype on expression on any of the homoeologs (ANOVA, p-value > 0.05, **Figure 1d-e**). This was further confirmed by the RNA-sequencing results (FDR adjusted p-value > 0.05, **Supplementary Figure 2a-b**). Overall, we found no evidence for widespread active transcriptional compensation between homoeologs in hexaploid wheat.

Next, we investigated whether active transcriptional compensation takes place in tetraploid wheat. As each homoeolog group consists of only two homoeologs (rather than three), a loss-of-function mutation in one homoeolog would result in a 50% reduction in functional transcript, compared to 33% in hexaploid wheat. Thus, we hypothesised that active transcriptional compensation would be more likely to occur in tetraploid wheat. From six EMS-mutagenised TILLING lines (cv. Kronos) (K2619, K2864, K3239, K0427, K4533, and K0774) a total of 101 unique homoeolog groups containing a PTC mutation were identified. In line with the results from hexaploid wheat, between 8.3% (in K0427) and 32.4% (in K2864) of the homoeologs with a PTC mutation were down-regulated in the mutagenised lines compared to WT (FDR adjusted p-value < 0.05, **Figure 2a**). We found three homoeolog groups (3.0%) affected by a PTC mutation in which the non-mutated homoeolog was up-regulated (FDR adjusted p-value < 0.05, **Figure 2a, Supplementary Table 1**). The proportion of homoeolog groups with up-regulation of the non-mutated homoeolog was not significantly different between those affected by a PTC mutation compared to those affected by a synonymous mutation (1.5% of homoeolog groups with a synonymous mutation, χ^2^ test p-value = 0.234, **Figure 2b**).

**Figure 2.**
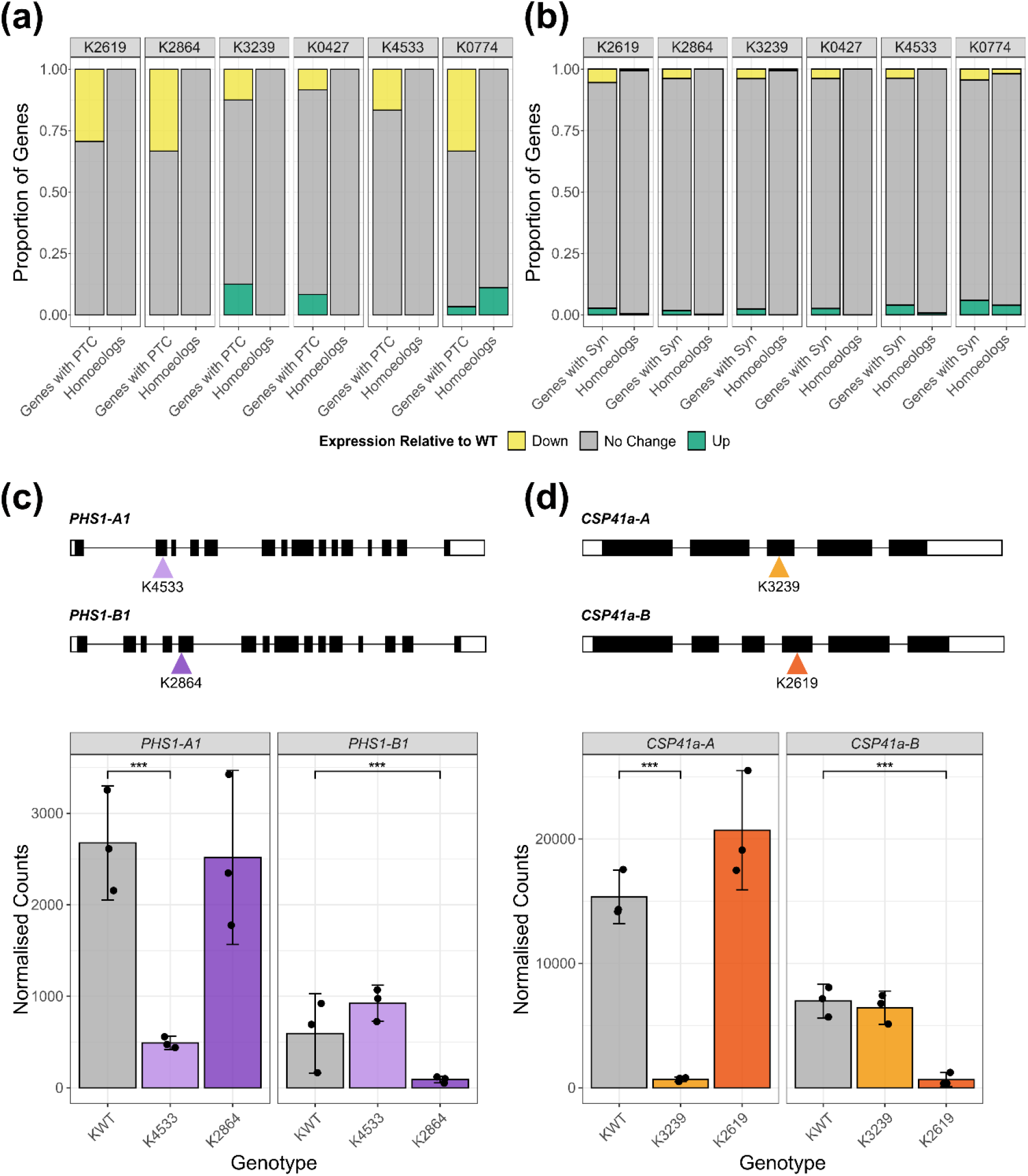
Absence of active transcriptional compensation between homoeologs in response to premature termination codon (PTC) mutations in tetraploid wheat. (a) Expression of genes with a PTC mutation (left bar) and their homoeologs (right bar) in each of the mutagenised Kronos lines. (b) Expression of genes with a synonymous mutation (left bar) and their homoeologs (right bar) in each of the mutagenised lines. In (a) and (b) the proportion of genes that are down-regulated regulated (FDR adjusted p-value < 0.05) is shown in yellow, up-regulated (FDR adjusted p-value < 0.05) in green, and no change in expression in grey, all relative to Kronos WT. (c) The location of PTC mutations in *PHS1-A1* and *PHS1-B1* in K4533 and K2864 respectively and the expression of *PHS1-A1* and *PHS1-B1* in Kronos WT (KWT) (grey), K4533 (light purple), and K2864 (dark purple). (d) The location of PTC mutations in *CSP41a-A* and *CSP41a-B* in K3239 and K2619 lines respectively and the expression of *CSP41a-A* and *CSP41a-B* in Kronos WT (KWT) (grey), K3239 (light orange), and K2619 (dark orange). In (c) and (d) in the gene schematics, black rectangles represent exons, white rectangles represent untranslated regions, and lines represent introns. Error bars represent the 95% confidence interval around the mean, estimated as 1.96 x standard error. Asterisks show significant differences in gene expression between gene expression; ***: p < 0.001 (DESeq2 p-value with FDR adjustment).

Despite the lack of widespread active transcriptional compensation between homoeologs in hexaploid and tetraploid wheat, we hypothesised that active transcriptional compensation may be responsible for the functional redundancy observed within particular homoeolog groups. The *PHS1* homoeolog group, encoding the plastidial α-glucan phosphorylase, has been characterised as functionally redundant by Kamble *et al*. (2023) – the *phs1-1* double mutant in tetraploid wheat has altered starch granule morphology (it has a decreased proportion of small starch granules and larger B-type granules), while neither the *phs1-A1* single mutant nor the *phs1-B1* single mutant has altered granule morphology. In contrast, the *CSP41a* homoeolog group, encoding a chloroplast RNA-binding protein, is non-redundant in tetraploid wheat – mutation of the A homoeolog alone (in the *csp41-A* single mutant) is sufficient to increase resistance to yellow rust (Corredor-Moreno *et al*., 2022). We used mutants in each of the homoeologs of *PHS1* and *CSP41a* to investigate whether the level of redundancy between homoeologs affected the degree of active transcriptional compensation observed. For both homoeologs of *PHS1*, the presence of a PTC mutation was associated with down-regulation of the affected homoeolog (for *PHS1-A1*, PTC mutation in K4533, fold change = 0.18, FDR adjusted p-value = 7.8×10^−19^; for *PHS1-B1*, PTC mutation in K2864, fold change = 0.15, FDR adjusted p-value = 4.3×10^−7^, **Figure 2c**). However, there was no up-regulation of the non-mutated homoeolog in either case (FDR adjusted p-value > 0.05, **Figure 2c**). Similarly, for *CSP41a*, the presence of a PTC mutation resulted in down-regulation of the affected homoeolog (for *CSP41a-A*, PTC mutation in K3239, fold change = 0.045, FDR adjusted p-value = 1.9×10^−61^; for *CSP41a-B*, PTC mutation in K2619, fold change = 0.093, FDR adjusted p-value = 7.6×10^−15^, **Figure 2d**). No up-regulation of the non-mutated homoeolog was observed in either case (FDR adjusted p-value > 0.05, **Figure 2d**). This indicates that active transcriptional compensation is not necessary for functional redundancy between homoeologs.

## Discussion

We have shown, for 188 homoeolog groups in hexaploid across two independent experiments and 101 homoeolog groups in tetraploid wheat, that active transcriptional compensation occurs at a low frequency – only 2.5-3.3% of PTC mutations were associated with up-regulation of the homoeologous genes across three independent datasets. These proportions were not significantly different from the proportion of synonymous mutations associated with homoeolog up-regulation, suggesting that apparent compensatory up-regulation of gene expression is not necessarily linked to the functional impact of the mutation (**Figure 1-2, Supplementary Figure 1**).

Mechanisms to compensate paralog loss-of-function have previously been shown to be non-universal - a similar study in yeast (*Saccharomyces cerevisiae*) found increased levels of approximately 11% of proteins when their paralogs were deleted (DeLuna *et al*., 2010). DeLuna *et al*. (2010) also showed that compensatory up-regulation occurred almost exclusively for proteins with essential functions, suggesting that up-regulation is induced in response to a physiological deficit, rather than as a direct response to the mutation. Such ‘needs-based’ responses could explain the up-regulation observed in the homoeolog group encoding a subunit of Respiratory Complex I (**Supplementary Table 1**; NADH dehydrogenase). However, this did not extend to any transcription factors, which are generally considered to be dosage-sensitive (Birchler & Veitia, 2010), and might therefore be expected to be transcriptionally compensated to maintain gene-dosage balance.

The low level of active transcriptional compensation observed in wheat is not in keeping with the functional redundancy that is often observed between homoeologs. Although almost all of the mutagenised genes in our study are functionally uncharacterised, the *PHS1* homoeologs in tetraploid wheat, previously shown to be functionally redundant, also did not show active transcriptional compensation (**Figure 2c**) (Kamble *et al*., 2023). It is possible that a different active compensatory mechanism facilitates functional redundancy between homoeologs – for example, Diss *et al*. (2014) propose translational up-regulation or changes in protein localisation as modes of compensation. Our study is also unlikely to capture cell-type or tissue-specific transcriptional compensation, as highlighted by Iohannes and Jackson (2023). Alternatively, there may be significant passive compensation between homoeologs, with two-thirds (in hexaploid wheat) or half (in tetraploid wheat) of the wild type levels of functional transcript being sufficient to maintain the wild-type phenotype.

Passive compensation between homoeologs in wheat is probable given that these gene duplications are relatively young and have had less time to diverge in expression levels – the hybridisation events occurred between 500,000 (forming A and B subgenomes) and 10,000 years ago (addition of the D subgenome) (El Baidouri *et al*., 2017). Increased time since gene duplication is likely to result in hypofunctionalisation, as the paralogs acquire mutations potentially decreasing their expression, and compensatory drift, in which the expression of one paralog decreases over time and function is maintained by its highly expressed copy (Iohannes & Jackson, 2023). When hypofunctionalisation or compensatory drift have occurred, expression from one paralog is insufficient to maintain gene function and an active compensatory mechanism would be required for functional redundancy (Iohannes & Jackson, 2023). Accordingly, Cusack *et al*. (2021) found that age of duplication is associated with functional redundancy in *A. thaliana*, with duplicates formed during the α-WGD more likely to be functionally redundant than those that arose during the more ancient β- or γ-WGD events. Furthermore, the *HBEGF* paralogs exhibiting active transcriptional compensation in zebrafish are much older than homoeologs in wheat, having arisen during the fish specific genome duplication event approximately 320 million years ago (Vandepoele *et al*., 2004; Laisney *et al*., 2010; El-Brolosy *et al*., 2019), and are thus more likely to have experienced hypofunctionalisation/compensatory drift. Similarly, the *CLV3* paralog in the Solanaceae, *CLE9*, arose at least 30 million years ago and shows active transcriptional compensation (Rodriguez-Leal *et al*., 2019). Overall, this underscores active transcriptional compensation as an evolutionary innovation that is only favourable when passive compensation is insufficient (whether to maintain functional redundancy or to adequately control gene expression noise, per Pires and Conant (2016)), and is otherwise unfavourable due to the cost of excess mRNA synthesis (with excess mRNA synthesis being selected against, as shown by Hausser *et al*. (2019)).

Passive compensation between homoeologs suggests that targeting active transcriptional compensation mechanisms characterised in other species is unlikely to break functional redundancy in wheat and would not be a viable strategy for breeding. Rather, mechanisms to simultaneously introduce mutations in multiple/all homoeologs, such as gene editing, or to reduce the expression of multiple/all homoeologs, such as RNAi, are advantageous to tackle functional redundancy.

## Materials & Methods

### Plant materials

Target genes were selected based on the availability of multiple EMS-mutagenised hexaploid wheat (*Triticum aestivum* cv. *Cadenza*) lines (Krasileva *et al*., 2017) with premature termination codon (PTC) mutations either early or late within the coding sequence of the same gene. Further screening of publicly available gene expression data (Borrill *et al*., 2016; Ramírez-González *et al*., 2018) was used to select genes with high expression in seedling leaf tissues. Based on these conditions, the genes *TraesCS5A02G119300* and *TraesCS6A02G155200* were chosen. Selected TILLING lines with PTC mutations in these genes are summarised below:

**Table 1:** Target genes and corresponding TILLING line mutations in Cadenza.

| Target Gene | TILLING Line | SNP | Amino Acid Change | Location Within Gene |
| --- | --- | --- | --- | --- |
| *TraesCS5A02G119300* | Cadenza1015 (C1015) | C/T | W/* | 2<sup>nd</sup> exon (of 9) |
|  | Cadenza1704 (C1704) | G/A | Q/* | 8<sup>th</sup> exon (of 9) |
| *TraesCS6A02G155200* | Cadenza0895 (C0895) | G/A | W/* | 1<sup>st</sup> exon (of 2) |
|  | Cadenza0604 (C0604) | G/A | W/* | 2<sup>nd</sup> exon (of 2) |

Single seed descent (SSD) lines were produced from the existing Cadenza TILLING M_5_ seed bulk for the selected lines (C0604, C0895, C1015, C1704) (Krasileva *et al*., 2017). For each line, a single M_6_ seed was sown and selfed in greenhouse conditions. The resulting M_7_ generation was grown in the field (Norwich, UK, 2017-2018) under standard agronomic practises in 1m^2^ plots to bulk seeds. The M_8_ seeds were sown for the RNA-sequencing experiment. The C0604, C0895, C1015, and C1704 SSD lines are available from the Germplasm Resources Unit, UK (www.seedstor.ac.uk).

For tetraploid wheat, target genes were selected from previously characterised redundant and non-redundant genes. Targets were chosen based on the availability of EMS-mutagenised tetraploid wheat (*Triticum turgidum* ssp. *durum* cv. *Kronos*) with PTC mutations in the coding sequence (Krasileva *et al*., 2017) and high expression in seedling leaf tissues. Selected TILLING lines with PTC mutations in these genes are summarised below:

**Table 2:** Target genes and corresponding TILLING line mutations in Kronos.

| Target Gene | Homoeolog | TILLING Line | SNP | Amino Acid Change | Location Within Gene |
| --- | --- | --- | --- | --- | --- |
| *PHS1* | *TraesCS5A02G395200* | Kronos4533 (K4533) | C/T | W/* | 2<sup>nd</sup> exon (of 15) |
|  | *TraesCS5B02G400000* | Kronos2864 (K2864) | C/T | W/* | 4<sup>th</sup> exon (of 15) |
| *CSP41a* | *TraesCS6A02G025700* | Kronos3239 (K3239) | G/A | Q/* | 3<sup>rd</sup> exon (of 6) |
|  | *TraesCS6B02G036400* | Kronos2619 (K2619) | C/T | W/* | 4<sup>th</sup> exon (of 6) |

In addition to Kronos4533, Kronos2864, Kronos3239, and Kronos2619, the TILLING lines Kronos0427 and Kronos0774 were grown, to adjust for the lower mutation density in the Kronos TILLING lines. Seeds for the selected Kronos lines were obtained from the Germplasm Resources Unit, UK (www.seedstor.ac.uk).

### Plant growth conditions

Seeds were imbibed on damp filter paper at 4°C for two days and then moved to room temperature to promote germination for a further two days. Seeds were then sown into 96 cell trays containing John Innes F_2_ Starter + Grit (90% peat, 10% grit, 4 kg m^−3^ dolomitic limestone, 1.2 kg m^−3^ osmocote start). Seedlings were grown in glasshouses at the John Innes Centre (Norwich, UK) with supplementary lighting and heating to maintain a minimum of 16 hours light, with 16°C day and 14°C night.

### DNA extraction & KASP genotyping

For Kronos, the TILLING lines were genotyped to ensure that they were homozygous for the PTC mutation of interest. DNA was extracted from seedling leaf tissue from two-week-old plants following the protocol from www.wheat-training.com. The KASP primers used for genotyping are shown in **Supplementary Table 2**. Genotyping was carried out using PACE mix (3CR Bioscience), according to the manufacturer’s instructions. Genotype calls were made using the Kluster-Caller software (v3.4.1.36).

### RNA extraction & RT-qPCR

Seedling leaf samples were taken from two-week-old plants (TILLING lines), frozen in liquid nitrogen, and stored at −70°C. RNA was extracted using the RNeasy Plant Mini Kit (Qiagen). Genomic DNA was digested using the RQ1 RNase-free DNase (Promega). cDNA synthesis was carried out using M-MLV reverse transcriptase (Invitrogen), according to the manufacturer’s instructions. Quantitative PCR (qPCR) was carried out in a LightCycler 480 (Roche) using the LightCycler 480 SYBR Green I Master (Roche) with the following cycling conditions: initial start of 95°C for 5 minutes; 45 cycles of 10 s at 95°C, 15 s at 60°C, 30 s at 72°C and reading for 1 s at 78°C. qPCR primers are shown in **Supplementary Table 2**. Relative expression was determined by calculating the Pfaffl ratio for each transcript relative to the GAPDH transcript. The effect of genotype on relative expression was tested using a linear model and ANOVA in R (v4.4.1).

### RNA extraction for RNA-seq & analysis

Samples were taken from the second leaf of two-week-old seedlings, frozen in liquid nitrogen and stored at −70°C. For the Cadenza samples, RNA was extracted using the Spectrum Plant Total RNA kit (Sigma) from three biological replicates for each TILLING line. For the Kronos samples, RNA was extracted using a Trizol-Chloroform method and cleaned with the RNA Clean & Concentrator Kit (Zymo Research). Library preparation and sequencing were carried out by Novogene (Cambridge, UK), producing 150 bp paired end reads. Reads were pseudoaligned to the IWGSC Chinese Spring v1.1 transcriptome (IWGSC *et al*., 2018) and quantified using kallisto (v0.46.1) (Bray *et al*., 2016). For the Kronos samples, reads were pseudoaligned to the Chinese Spring transcriptome with transcripts from the D subgenome removed. An average of 28,429,382 reads per sample (81% of the total reads) were pseudoaligned for the Cadenza TILLING lines and an average of 27,411,192 reads per sample (74% of the total reads) were pseudoaligned for the Kronos TILLING lines. Abundance files from kallisto were imported into R (v4.4.1) using Tximport (Soneson *et al*., 2015). Differentially expressed genes between the TILLING lines and WT were identified using DESeq2 (Love et al., 2014), applying a threshold of adjusted p-value < 0.05. Up-regulated genes were classified as those with fold change > 1, and down-regulated genes as those with fold change < 1.

To identify variants present in the TILLING lines, the raw reads were trimmed with trimmomatic (v0.39) (parameters: ILLUMINACLIP:TruSeq3-PE.fa:2:30:10 LEADING:3 TRAILING:3 SLIDINGWINDOW:4:15 MINLEN:80) (Bolger *et al*., 2014) and mapped to the IWGSC Chinese Spring v1.0 reference sequence (IWGSC *et al*., 2018) using HISAT2 (v2.1.0) (Pertea *et al*., 2016) and SAMtools (v1.12) (Danecek *et al*., 2021). Freebayes (v1.2.0) was used for variant calling (parameters: -F 0.87 --min-coverage 10 --use-best-n-alleles 2) (Garrison & Marth, 2012) and variant effect prediction was carried out using VEP (v91.3) (McLaren *et al*., 2016). Variants were filtered to keep only those predicted to cause a premature termination codon or synonymous mutation and were homozygous. Further filtering was carried out to identify variants in homoeologous genes (using homoeolog groups defined in Evans *et al*. (2022)), and those present in one of the TILLING lines, but not WT to account for cultivar specific SNPs. The lists of differentially expressed genes were then used to identify changes in expression in the genes affected by a PTC/synonymous mutation and their homoeologs.

The same analysis was carried out on RNA-sequencing data from Xiong, HC *et al*. (2020), comparing two independent EMS-mutagenised lines (*dm3* and *dm4*) to the Jing411 WT control.

## Acknowledgements

We thank Marek Glombik for advice on RNA-sequencing analysis, and Cristobal Uauy for feedback during the project. This research was supported by NBI Research Computing through HPC resources. This work was supported by the UK Biotechnology and Biological Science Research Council (BBSRC) through grant BB/T013524/2 and the Institute Strategic Programmes Delivering Sustainable Wheat (DSW) (BB/X011003/1) and Building Robustness in Crops (BRiC) (BB/X01102X/1). DD was funded through a BBSRC Doctoral Training Partnership (BB/T008717/1).

## Author Contributions

PB conceived the study. DD and PB designed the research. JS generated the Cadenza SSD lines. DD and EK performed experiments. DD carried out analysis of RNA-sequencing data. DD created the figures. DD and PB wrote the manuscript with input from EK and JS. All authors have read and approved the manuscript.

## Data Availability

The Cadenza SSD lines (C0604, C0895, C1015, and C1704) are available from the Germplasm Resources Unit (Norwich, United Kingdom; www.seedstor.ac.uk) through the Deposited Published Research Material Collection.

The Kronos EMS TILLING lines can be ordered from the Germplasm Resources Unit (Norwich, United Kingdom; www.seedstor.ac.uk) under accession names Kronos0427, Kronos0774, Kronos2619, Kronos2864, Kronos3239, and Kronos4533.

Raw reads from RNA-sequencing have been deposited in the European Nucleotide Archive under project PRJEB89501.

Scripts used for analysis of the RNA-sequencing data can be found on GitHub at www.github.com/Borrill-Lab/Transcriptional_Compensation.

## Supporting Information

**Supplementary Figure 1**. Absence of widespread active transcriptional compensation in an independent RNA-seq dataset of EMS-mutagenised hexaploid wheat from Xiong, HC *et al*. (2020).

**Supplementary Figure 2**. PTC location does not affect whether transcriptional compensation occurs between homoeologs.

**Supplementary Table 1**. Homoeolog groups in which at least one non-mutated homoeolog is up-regulated relative to the WT control.

**Supplementary Table 2**. Sequences of all primers used in this study.

## References

Abe F, Haque E, Hisano H, Tanaka T, Kamiya Y, Mikami M, Kawaura K, Endo M, Onishi K, Hayashi T, Sato K. 2019. Genome-edited triple-recessive mutation alters seed dormancy in wheat. Cell Rep 28(5): 1362–1369.

Bailon-Zambrano R, Sucharov J, Mumme-Monheit A, Murry M, Stenzel A, Pulvino AT, Mitchell JM, Colborn KL, Nichols JT. 2022. Variable paralog expression underlies phenotype variation. Elife 11: e79247.

Birchler JA, Veitia RA. 2010. The gene balance hypothesis: implications for gene regulation, quantitative traits and evolution. New Phytologist 186(1): 54–62.

Birchler JA, Yang H. 2022. The multiple fates of gene duplications: deletion, hypofunctionalization, subfunctionalization, neofunctionalization, dosage balance constraints, and neutral variation. Plant Cell 34(7): 2466–2474.

Bolger AM, Lohse M, Usadel B. 2014. Trimmomatic: a flexible trimmer for Illumina sequence data. Bioinformatics 30(15): 2114–2120.

Borrill P, Adamski N, Uauy C. 2015. Genomics as the key to unlocking the polyploid potential of wheat. New Phytologist 208(4): 1008–1022.

Borrill P, Ramirez-Gonzalez R, Uauy C. 2016. expVIP: a customizable RNA-seq data analysis and visualization platform. Plant Physiol 170(4): 2172–2186.

Bray NL, Pimentel H, Melsted P, Pachter L. 2016. Near-optimal probabilistic RNA-seq quantification. Nature Biotechnology 34(5): 525–527.

Corredor-Moreno P, Badgami R, Jones S, Saunders DGO. 2022. Temporally coordinated expression of nuclear genes encoding chloroplast proteins in wheat promotes Puccinia striiformis f. sp. tritici infection. Commun Biol 5(1): 853.

Cusack SA, Wang PP, Lotreck SG, Moore BM, Meng FR, Conner JK, Krysan PJ, Lehti-Shiu MD, Shiu SH. 2021. Predictive models of genetic redundancy in Arabidopsis thaliana. Molecular Biology and Evolution 38(8): 3397–3414.

Danecek P, Bonfield JK, Liddle J, Marshall J, Ohan V, Pollard MO, Whitwham A, Keane T, McCarthy SA, Davies RM, Li H. 2021. Twelve years of SAMtools and BCFtools. Gigascience 10(2): giab008.

Degenhardt RF, Bonham-Smith PC. 2008. Transcript profiling demonstrates absence of dosage compensation in Arabidopsis following loss of a single RPL23a paralog. Planta 228(4): 627–640.

DeLuna A, Springer M, Kirschner MW, Kishony R. 2010. Need-based up-regulation of protein levels in response to deletion of their duplicate genes. Plos Biology 8(3): e1000347.

Diss G, Ascencio D, DeLuna A, Landry CR. 2014. Molecular mechanisms of paralogous compensation and the robustness of cellular networks. Journal of Experimental Zoology Part B-Molecular and Developmental Evolution 322(7): 488–499.

El Baidouri M, Murat F, Veyssiere M, Molinier M, Flores R, Burlot L, Alaux M, Quesneville H, Pont C, Salse J. 2017. Reconciling the evolutionary origin of bread wheat (Triticum aestivum). New Phytologist 213(3): 1477–1486.

El-Brolosy MA, Kontarakis Z, Rossi A, Kuenne C, Günther S, Fukuda N, Kikhi K, Boezio GLM, Takacs CM, Lai SL, et al. 2019. Genetic compensation triggered by mutant mRNA degradation. Nature 568(7751): 193–197.

Evans CEB, Arunkumar R, Borrill P. 2022. Transcription factor retention through multiple polyploidization steps in wheat. G3-Genes Genomes Genetics 12(8): jkac147.

Garrison E, Marth G. 2012. Haplotype-based variant detection from short-read sequencing. DOI: 10.48550/arXiv.1207.3907

Hausser J, Mayo A, Keren L, Alon U. 2019. Central dogma rates and the trade-off between precision and economy in gene expression. Nature Communications 10: 68.

Iohannes SD, Jackson D. 2023. Tackling redundancy: genetic mechanisms underlying paralog compensation in plants. New Phytologist 240(4): 1381–1389.

Iwgsc, Appels R, Eversole K, Stein N, Feuillet C, Keller B, Rogers J, Pozniak CJ, Choulet F, Distelfeld A, et al. 2018. Shifting the limits in wheat research and breeding using a fully annotated reference genome. Science 361(6403): eaar7191.

Kamble NU, Makhamadjonov F, Fahy B, Martins C, Saalbach G, Seung D. 2023. Initiation of B-type starch granules in wheat endosperm requires the plastidial ɑ-glucan phosphorylase PHS1. Plant Cell 35(11): 4091–4110.

Krasileva KV, Vasquez-Gross HA, Howell T, Bailey P, Paraiso F, Clissold L, Simmonds J, Ramirez-Gonzalez RH, Wang XD, Borrill P, et al. 2017. Uncovering hidden variation in polyploid wheat. Proceedings of the National Academy of Sciences of the United States of America 114(6): E913–E921.

Kwon CT, Tang LL, Wang XG, Gentile I, Hendelman A, Robitaille G, Van Eck J, Xu C, Lippman ZB. 2022. Dynamic evolution of small signalling peptide compensation in plant stem cell control. Nature Plants 8(4): 346–355.

Laisney JAGC, Braasch I, Walter RB, Meierjohann S, Schartl M. 2010. Lineage-specific co-evolution of the Egf receptor/ligand signaling system. Bmc Evolutionary Biology 10: 27.

Ma ZP, Zhu PP, Shi H, Guo LW, Zhang QH, Chen YN, Chen SM, Zhang Z, Peng JR, Chen J. 2019. PTC-bearing mRNA elicits a genetic compensation response via Upf3a and COMPASS components. Nature 568(7751): 259–263.

Matsuoka Y. 2011. Evolution of polyploid Triticum wheats under cultivation: the role of domestication, natural hybridization and allopolyploid speciation in their diversification. Plant and Cell Physiology 52(5): 750–764.

McLaren W, Gil L, Hunt SE, Riat HS, Ritchie GR, Thormann A, Flicek P, Cunningham F. 2016. The Ensembl Variant Effect Predictor. Genome Biology 17(1): 122.

Panchy N, Lehti-Shiu M, Shiu SH. 2016. Evolution of gene duplication in plants. Plant Physiology 171(4): 2294–2316.

Pertea M, Kim D, Pertea GM, Leek JT, Salzberg SL. 2016. Transcript-level expression analysis of RNA-seq experiments with HISAT, StringTie and Ballgown. Nat Protoc 11(9): 1650–1667.

Pires JC, Conant GC. 2016. Robust yet fragile: expression noise, protein misfolding, and gene dosage in the evolution of genomes. Annual Review of Genetics 50: 113–131.

Ramírez-González RH, Borrill P, Lang D, Harrington SA, Brinton J, Venturini L, Davey M, Jacobs J, van Ex F, Pasha A, et al. 2018. The transcriptional landscape of polyploid wheat. Science 361(6403): eaar6089.

Rodriguez-Leal D, Xu C, Kwon CT, Soyars C, Demesa-Arevalo E, Man J, Liu L, Lemmon ZH, Jones DS, Van Eck J, et al. 2019. Evolution of buffering in a genetic circuit controlling plant stem cell proliferation. Nature Genetics 51(5): 786–792.

Schreiber AW, Hayden MJ, Forrest KL, Kong SL, Langridge P, Baumann U. 2012. Transcriptome-scale homoeolog-specific transcript assemblies of bread wheat. Bmc Genomics 13: 492.

Serobyan V, Kontarakis Z, El-Brolosy MA, Welker JM, Tolstenkov O, Saadeldein AM, Retzer N, Gottschalk A, Wehman AM, Stainier DYR. 2020. Transcriptional adaptation in Caenorhabditis elegans. Elife 9: e50014.

Singh M, Kumar M, Thilges K, Cho MJ, Cigan AM. 2017. MS26/CYP704B is required for anther and pollen wall development in bread wheat (Triticum aestivum L.) and combining mutations in all three homeologs causes male sterility. PLoS One 12(5): e0177632.

Soneson C, Love MI, Robinson MD. 2015. Differential analyses for RNA-seq: transcript-level estimates improve gene-level inferences. F1000Res 4: 1521.

Uauy C, Wulff BBH, Dubcovsky J. 2017. Combining traditional mutagenesis with new high-throughput sequencing and genome editing to reveal hidden variation in polyploid wheat. Annual Review of Genetics, Vol 51 51: 435–454.

Vandepoele K, De Vos W, Taylor JS, Meyer A, Van de Peer Y. 2004. Major events in the genome evolution of vertebrates: paranome age and size differ considerably between ray-finned fishes and land vertebrates. Proceedings of the National Academy of Sciences of the United States of America 101(6): 1638–1643.

Xiong HC, Zhou CY, Guo HJ, Xie YD, Zhao LS, Gu JY, Zhao SR, Ding YP, Liu LX. 2020. Transcriptome sequencing reveals hotspot mutation regions and dwarfing mechanisms in wheat mutants induced by γ-ray irradiation and EMS. Journal of Radiation Research 61(1): 44–57.

Xiong W, Chen XZ, Zhu CX, Zhang JC, Lan T, Liu L, Mo BX, Chen XM. 2020. Arabidopsis paralogous genes RPL23aA and RPL23aB encode functionally equivalent proteins. Bmc Plant Biology 20(1): 463.

