## Supplementary Figures and Tables for "Homoeolog expression in polyploid wheat mutants shows limited transcriptional compensation"

Delfi Dorussen[1], Emilie Knight[1], James Simmonds[1], Philippa Borrill[1*]

[1] Department of Crop Genetics, John Innes Centre, Norwich Research Park, Norwich, NR4 7UH, United Kingdom

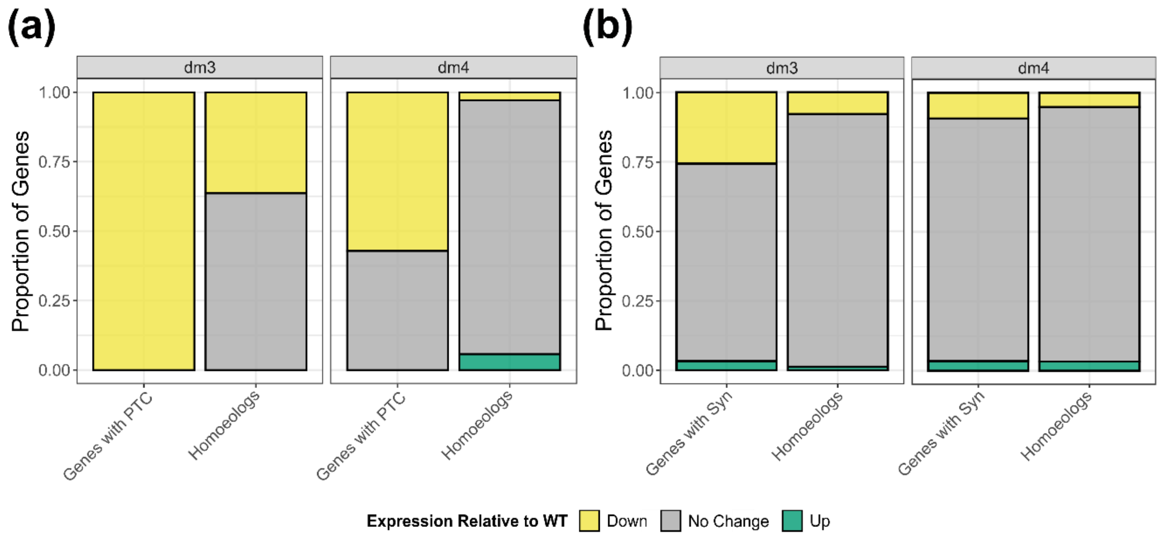


**Supplementary Figure 1. Absence of widespread active transcriptional compensation in an independent RNA-seq dataset of EMS-mutagenised hexaploid wheat from Xiong, HC *et al.* (2020).** (a) Expression of genes with a premature termination codon (PTC) mutation (left bar) and their homoeologs (right bar) in each of the EMS-mutagenised lines (*dm3* and *dm4*). (b) Expression of genes with a synonymous mutation (left bar) and their homoeologs (right bar) in each of the EMS-mutagenised lines (dm3 and dm4). In (a) and (b) the proportion of genes that are down-regulated (FDR adjusted p-value < 0.05) is shown in yellow, up-regulated (FDR adjusted p-value < 0.05) in green, and no change in expression in grey, all relative to Jing411 WT.

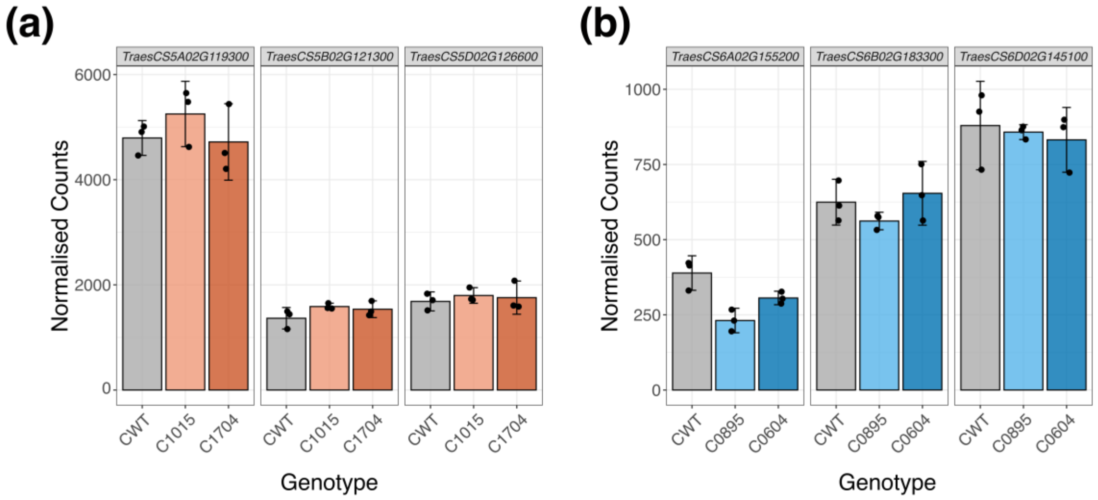


**Supplementary Figure 2. PTC location does not affect whether active transcriptional compensation occurs between homoeologs.** (a) The DESeq2 normalised counts for *TraesCS5A02G119300* and its homoeologs (*TraesCS5B02G121300* and *TraesCS5D02G126600*) in Cadenza WT (CWT) (grey), C1015 (light orange), and C1704 (dark orange). (b) The DESeq2 normalised counts for *TraesCS6A02G155200* and its homoeologs (*TraesCS6B02G183300* and *TraesCS6D02G145100*) in Cadenza WT (CWT) (grey), C0895 (light blue), and C0604 (dark blue). Error bars represent the 95% confidence interval around the mean, estimated as 1.96 x standard error.

**Supplementary Tables**

**Supplementary Table 1. Homoeolog groups in which at least one non-mutated homoeolog is up-regulated relative to the WT control.**

| Mutant Line/ Source | Annotation of Homoeolog Group | Homoeologs | PTC Mutation | Fold Change | DESeq2 Adjusted p-value | Expression Relative to WT |
|---|---|---|---|---|---|---|
| C0604 | MFS transporter (IPR036259); proton-dependent oligopeptide transporter (IPR000109) | *TraesCS1A02G269600* | No | 1.98 | 0.000317 | Up |
| | | *TraesCS1B02G280100* | No | 1.47 | 0.134 | - |
| | | *TraesCS1D02G269700* | Yes (Exon 4/4; AA 529/576) | 1.74 | 0.0201 | Up |
| C0895 | Meiosis regulator and mRNA stability factor 1 (IPR024768) | *TraesCS2A02G150200* | No | 3.48 | 0.0125 | Up |
| | | *TraesCS2B02G175400* | Yes (Exon 2/2; AA 152/632) | 0.50 | 0.540 | - |
| | | *TraesCS2D02G155300* | No | 0.64 | 0.559 | - |
| C1015 | MFS transporter (IPR036259); sugar phosphate transporter (IPR000849) | *TraesCS7A02G300800* | No | 1.76 | 0.0349 | Up |
| | | *TraesCS7B02G201500* | No | 1.53 | 0.0609 | - |
| | | *TraesCS7D02G296500* | Yes (Exon 1/2; AA 272/559) | 1.08 | 0.912 | - |
| | Serine/ threonine protein kinase (IPR011009; IPR008271) | *TraesCS7A02G544000* | Yes (Exon 1/1; AA 26/413) | 2.16 | 0.342 | - |
| | | *TraesCS7B02G466300* | No | 1.03 | 0.970 | - |
| | | *TraesCS7D02G530500* | No | 4.31 | 0.000108 | Up |
| dm4 (Xiong HC *et al.*, 2020) | NADH dehydrogenase [ubiquinone] (complex I), alpha subcomplex subunit 1 (IPR017384) | *TraesCS1A02G302900* | Yes (Exon 1/3; AA 8/66) | 0.11 | $7.31 \times 10^{-82}$ | Down |
| | | *TraesCS1B02G313200* | No | 1.45 | 0.0130 | Up |
| | | *TraesCS1D02G302300* | No | 1.45 | 0.00348 | Up |
| K0774 | Cytokinin dehydrogenase (IPR016170; IPR015345) | *TraesCS1A02G159600* | No | 2.65 | 0.00183 | Up |
| | | *TraesCS1B02G176000* | Yes (Exon 5/6; AA 338/522) | 2.07 | 0.0329 | Up |
| | ATP-dependent Clp protease/ Chaperone ClpA/ClpB (IPR050130) | *TraesCS2A02G291100* | Yes (Exon 5/12; AA 420/946) | 1.59 | 0.162 | - |
| | | *TraesCS2B02G307500* | No | 1.76 | 0.00771 | Up |
| | Squalene/ phytoene synthase (IPR002060) | *TraesCS5A02G020900* | Yes (Exon 1/6; AA 27/396) | 0.63 | 0.189 | - |
| | | *TraesCS5B02G017900* | No | 2.64 | 0.00860 | Up |

**Supplementary Table 2. Sequences of all primers used in this study.** For KASP genotyping, the VIC tail (GAAGGTCGGAGTCAACGGATT) was added to the 5' end of the WT allele primers, and the FAM tail (GAAGGTGACCAAGTTCATGCT) was added to the 5' end of the mutant allele primers.

| **Name of Primer** | **Sequence (5'-3')** | **Purpose** |
|---|---|---|
| Gene4_F1_Aspe | CTACCGATTCTTGGAGGAAGG | qPCR; forwards primer for A homoeolog of *TraesCS5A02G119300* |
| Gene4_F2_Bspe | ATGGAGCTTCTGTCTCACATGT | qPCR; forwards primer for B homoeolog of *TraesCS5A02G119300* |
| Gene4_F1_Dspe | CTACCGATTCTTGGAGGAACA | qPCR; forwards primer for D homoeolog of *TraesCS5A02G119300* |
| Gene4_R1 | CCCATTCAGGTTCTGCTTTCT | qPCR; reverse primer for A and D homoeologs of *TraesCS5A02G119300* |
| Gene4_R2 | TGATTCTGGATGTGCTTCCT | qPCR; reverse primer for B homoeolog of *TraesCS5A02G119300* |
| Gene3_F1_Aspe | GCTGCTGGAGAAGCCTGAA | qPCR; forwards primer for A homoeolog of *TraesCS6A02G155200* |
| Gene3_F1_Bspe | GCTGCTGGAGAAGCCTGAG | qPCR; forwards primer for B homoeolog of *TraesCS6A02G155200* |
| Gene3_F1_Dspe | GCTGCTGGAGAAGCCTGAC | qPCR; forwards primer for D homoeolog of *TraesCS6A02G155200* |
| Gene3_R1 | ATATTTGGGTGCCTTAGGGTAGA | qPCR; reverse primer for A, B and D homoeologs of *TraesCS6A02G155200* |
| TaGAPDH_F | TTAGACTTGCGAAGCCAGCA | qPCR; forwards primer for GAPDH control |
| TaGAPDH_R | AAATGCCCTTGAGGTTTCCC | qPCR; reverse primer for GAPDH control |
| PHS1-A_K4533_WT | TTTTTGATTCTCTGATATTGAATTG | KASP genotyping; WT primer for K4533 (*phs1-a1* mutant); from Kamble *et al*. (2023) |
| PHS1-A_K4533_Mut | TTTTTGATTCTCTGATATTGAATTA | KASP genotyping; mutant primer for K4533 (*phs1-a1* mutant); from Kamble *et al*. (2023) |
| PHS1-A_K4533_Com | AAGGTATGAGATAAGGCTGAAGA | KASP genotyping; common primer for K4533 (*phs1-a1* mutant); from Kamble *et al*. (2023) |
| PHS1-B_K2864_WT | AGAGACATCATTTCTTACGATCTCC | KASP genotyping; WT primer for K2864 (*phs1-b1* mutant); from Kamble *et al*. (2023) |

| | | |
|---|---|---|
| PHS1-B_K2864_Mut | AGAGACATCATTTCTTACGATCTCT | KASP genotyping; mutant primer for K2864 (*phs1-b1* mutant); from Kamble *et al*. (2023) |
| PHS1-B_K2864_Com | ACCTGCATGTTAGCTTCTTTTCT | KASP genotyping; common primer for K2864 (*phs1-b1* mutant); from Kamble *et al*. (2023) |
| CSP41a-A_K3239_WT | GCTGCTGATGAACAGGAACTG | KASP genotyping; WT primer for K3239 (*csp41-a* mutant); from Corredor-Moreno *et al*. (2022) |
| CSP41a-A_K3239_Mut | GCTGCTGATGAACAGGAACTA | KASP genotyping; mutant primer for K3239 (*csp41-a* mutant); from Corredor-Moreno *et al*. (2022) |
| CSP41a-A_K3239_Com | AGTTTTGATGTTTGGTTGCGT | KASP genotyping; common primer for K3239 (*csp41-a* mutant); from Corredor-Moreno *et al*. (2022) |
| CSP41a-B_K2619_WT_1 | GCGGAGTTCGGCAGCTGG | KASP genotyping; WT primer for K2619 (*csp41-b* mutant) |
| CSP41a-B_K2619_Mut_1 | GCGGAGTTCGGCAGCTGA | KASP genotyping; mutant primer for K2619 (*csp41-b* mutant) |
| CSP41a-B_K2619_Com_1a | CTCGCAGTCCTTGTTGTTGC | KASP genotyping; common primer for K2619 (*csp41-b* mutant) |
